# The *C. elegans* CHP1 homolog, *pbo-1*, functions in innate immunity by regulating the pH of the intestinal lumen

**DOI:** 10.1101/424382

**Authors:** Saida Benomar, Aaron Bender, Blake R. Peterson, Josephine R. Chandler, Brian D. Ackley

**Author notes:** Co-corresponding authors: Josephine R. Chandler, Department of Molecular Biosciences, 1200 Sunnyside Ave, Lawrence, KS 66045,; Brian D. Ackley, Department of Molecular Biosciences, 1200 Sunnyside Ave, Lawrence, KS 66045.

## Abstract

*Caenorhabditis elegans* are soil-dwelling nematodes and models for understanding innate immunity and infection. Previous work has described a regularly-timed pH change in the intestine of *Caenorhabditis elegans* called the pH wave. To characterize this wave and its function in the worm, we developed a novel fluorescent dye (KR35) that accumulates in the intestine and sensitively responds to dynamic changes in pH. Here, we use KR35 to show that mutations in the Ca^2+^-binding protein, PBO-1 abrogate the pH wave, causing the anterior intestine to be constantly acidic. Surprisingly, *pbo-1* mutants were also more susceptible to infection by several bacterial pathogens. We could suppress pathogen susceptibility in *pbo-1* mutants by treating the animals with pH-buffering bicarbonate, suggesting the pathogen susceptibility is a function of the acidity of the intestinal pH. Furthermore, we use KR35 to show that pathogens completely neutralize the pH in the intestine of wild type, but not *pbo-1* mutants. *C. elegans* is known to increase production of reactive oxygen species (ROS), such as H_2_O_2,_ in response to pathogens, which is an important component of pathogen defense. We show that *pbo-1* mutants exhibited decreased H_2_O_2_ in response to pathogens, which could also be partially restored in *pbo-1* animals treated with bicarbonate. Ultimately, our results support a model whereby *pbo-1* functions during infection to permit pH changes in the intestine that are important for fighting pathogens.

**Author Summary:** Innate immunity is critical for host defense against pathogens. However, questions remain about how the host senses and responds to pathogen invasion. Using a pH-sensitive fluorescent dye and a *Caenorhabditis elegans* pathogen infection model we show that pathogens induce changes in pH of the worm intestine. We also show that intestinal pH directly affects production of reactive oxygen species (e.g. H_2_O_2_) important for pathogen defense. Our results show that pH regulation is an important component of the innate immune response to pathogens.

## Introduction

As increased antibiotic resistance is being observed in clinical settings, bacterial infections are becoming a crisis-level global health burden. Host barriers to infection can be protective against dangerous infections in the absence of antibiotics, and new discoveries about the mechanisms of host protection might pave the way for new therapeutics to treat disease. Host barriers to infection can be physical (*e.g*. an exoskeleton or epidermal layer), or chemical (*e.g*. shed-able mucosa that adhere to pathogens) and genetic (*e.g*. innate and/or adaptive immunity).

One particularly effective mechanism of host protection is the production of reactive oxygen species (ROS), *e.g*. superoxide (O_2_^−^) which can be converted to H_2_O_2_ [1]. Over the past 15–20 years H2O_2_ has emerged as a critical signaling molecule in several contexts, including infection and wound repair [2-4]. In the vertebrate innate immune response, the enzymes NADPH oxidase and superoxide dismutase (SOD) convert O_2_ to O_2_^-^ and H_2_O_2_ [5, 6]. H_2_O_2_ can be produced intracellularly and exported via aquaporin channels or produced in the extracellular space of cells responding to bacterial infection. Because ROS can also be self-destructive, it is no surprise that the production of ROS is tightly regulated in response to pathogens [7]. However, the identities of infection-dependent signals and regulatory responses that lead to ROS production remain incomplete.

Many discoveries about the role of ROS and other innate immune responses in organismal defenses to pathogens have come from genetically-tractable model systems where infection can be presented in a controlled manner. For example, the nematode *Caenorhabditis elegans* has served as an excellent system to understand the mechanisms that underlie organismal responses to pathogens and wounding [8-10]. Portions of the ROS production pathways are largely conserved in *C. elegans* and vertebrate animals, including humans. For example *itr-1*, the inositol trisphosphate receptor is required for intracellular calcium release in response to infection [1], an aquaporin, *aqp-1*, is required for the full anti-pathogen response [11], and a dual oxidase/peroxidase, *bli-3* and a secreted peroxidase, *skpo-1*, are important for the defense against pathogens [12, 13]. Importantly, these pathways are simplified in *C. elegans* with minimal redundancy. Taken together, the evolutionary conservation of organismal response to infection indicates that we can use *C. elegans* to better understand vertebrate response to bacterial pathogens.

This study is focused on PBO-1, a protein we find to be key in the response to pathogen infection. PBO-1 was first identified because of its contribution to the posterior body contraction (Pbo) phenotype [14]. PBO-1 is an EF-hand protein orthologous to human CHP1 (Calcineurin B homologous protein 1, also called P22) [15]. It is thought that PBO-1 links the release of intracellular calcium to the activity and/or trafficking of Na^+^/H^+^-exchangers (NHE or NHX), including NHX-2, NHX-6 and PBO-4 (also known as NHX-7) [16]. In *C. elegans*, PBO-4 is required for the release of protons from the basolateral membrane of the intestinal cell onto adjacent muscles, where they activate the proton-gated ion channel, PBO-5 [17].

In this study, we show that PBO-1 loss-of-function mutations fail to undergo oscillatory pH fluctuations in the intestine and have an intestinal pH that is more acidic than wild-type animals. Surprisingly, *pbo-1* mutants are more susceptible to pathogens than wild-type animals and show decreased production of ROS during infection. Our results also show that pathogen infections alter the intestinal pH in *C. elegans*, and that this regulation is at least partially dependent on PBO-1. We can suppress the acidic phenotype and pathogen susceptibility by removing PBO-4 from *pbo-1* mutant animals or by adding pH-neutralizing sodium bicarbonate to the *C. elegans* growth medium. Overall, we propose a model whereby infection by pathogens initiates a neutralization of the normally acidic pH in the intestine, which enables increased levels of ROS production to fight the pathogen.

## Results

### pbo-1 mutants have an abrogated pH wave in the intestine

Previously we developed a pH-responsive dye, KR35, and used it to monitor pH changes in the intestine of *C. elegans* [18]. KR35 is activated by acid in a range that is physiologically relevant to the *C. elegans* intestine, which maintains a pH gradient from anterior to posterior of ~5.5-3.5 [19]. When we feed wild-type (N2) animals KR35 we observe fluorescence in the posterior of the *C. elegans* intestine, indicating acidity in this region. We also observe a dynamic change in pH or an ‘acidic wave’, with a periodicity of ~50 seconds (Fig. 1, Supp. Video 1). During the oscillation, the posterior fluorescence transitions to the anterior region of the intestine near the pharyngeal-intestinal junction and stays there for about 3-7 seconds during a period we have termed the <u>M</u>aximum <u>A</u>nterior <u>T</u>ransition (MAT). The 50 second oscillations and the MAT are part of a process called the defecation motor program (DMP), whereby protons are transported across the basolateral membrane of the intestine by the Na^+^/H^+^ exchanger PBO-4, where they act as neurotransmitters that trigger muscle contractions required for elimination of waste [16, 17]. Using KR35 we previously showed that animals with mutations in *pbo-4* fail to produce the MAT during pH oscillations, although other aspects of the acidic wave were grossly normal such as the acidification of the posterior intestine prior to the MAT [18].

**Fig. 1.**
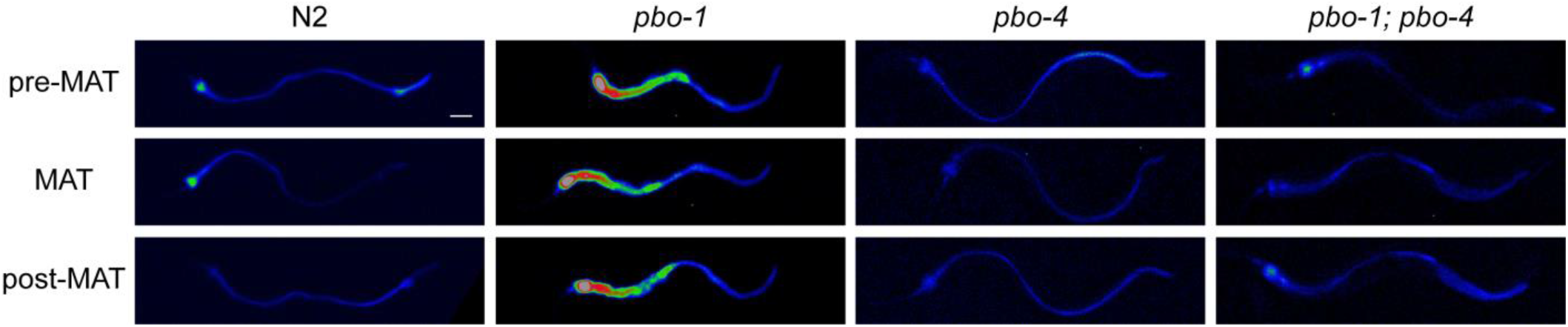
*pbo-1* mutations disrupt the intestinal acid wave. Fluorescence micrographs of *C. elegans* wild type or *pbo* mutants after feeding on the pH-sensitive probe KR35 for 30 min (10 M). Time-dependent images during the DMP extracted from video microscopy are shown. Fluorescence is rendered as a heat map of fluorescence intensity with red representing the most intense fluorescence (high acidity), and black the least intense fluorescence (low acidity). The head of the animal is on the left side of each image (scale bar = 50 μm).

During the defecation motor program (DMP), *pbo-4* mutants exhibit a <u>P</u>osterior <u>bo</u>dy contraction absent phenotype (Pbo) because of the defect in contracting posterior muscles during DMP [17]. To evaluate the relationship between the MAT and Pbo phenotype, we examined another Pbo mutant, *pbo-1*(*sa7*). We found that *pbo-1* mutants fed KR35 failed to demonstrate any pH oscillations and instead had an anterior intestine that was largely acidified relative to wild type (Fig. 1, Supp. Video 2). The posterior intestine of the *pbo-1* mutant was less acidic than wild type (Fig. 1), which likely explains the Pbo phenotype because proton transport from the posterior intestine onto the body wall muscle is required to stimulate muscle contraction during the normal DMP [17]. The anterior acidity in *pbo-1* mutants was the opposite of the *pbo-4* phenotype, where animals fail to acidify the anterior (Fig. 1, Supp. Video 3). Thus, we analyzed *pbo-1; pbo-4* double mutant animals using KR35. *pbo-1, pbo-4* double mutants had reduced KR35 fluorescence (more neutral intestinal pH) compared with *pbo-1* single mutants. Also, unlike *pbo-1* single mutants, the *pbo-1, pbo-4* double mutants exhibited pH oscillations, and protons were evacuated from the anterior during the MAT (Fig 1, Supp. Video 4). Our observation that the *pbo-1, pbo-4* double mutants are more similar to the *pbo-4* single mutants suggests the acidic phenotype of the *pbo-1* mutants is dependent on *pbo-4* function.

Our results with KR35 are consistent with the known role of PBO-1 in trafficking PBO-4 [20], and more generally of the role of PBO-1-like molecules being regulators of NHX activity [21]. Our results would suggest, at least in the anterior of the animal, that PBO-1 is a negative regulator of PBO-4. In this case, loss of PBO-1 function would aberrantly activate PBO-4, leading to a PBO-4-dependent accumulation of protons in the anterior. Removing *pbo-4* from the *pbo-1* mutant abrogates these effects, and somewhat normalizes the acidic phenotype of the *pbo-1* mutant (Fig. 1). *pbo-1; pbo-4* double mutants were not identical to the *pbo-4* single mutants, and these differences likely come from the function of other NHX proteins in the intestine [16], which are likely to be regulated by PBO-1.

### pbo-1 mutants are more susceptible to pathogens

Our results with KR35 support the conclusion that defects in the dynamic acidic wave can lead to the Pbo phenotype, and not the other way around. We hypothesized that the acidic wave may have other physiological consequences to the animals. For example, low intestinal pH could function to kill pathogens that entered the alimentary canal during feeding or begin digestion of food deposited in the intestine, analogous to the function of low pH in the stomach of vertebrates. In other words, perhaps the animals create a temporary region of very low pH to kill pathogens. To test this notion, we fed wild type and *pbo-1* mutants the opportunistic pathogen *Enterococcus faecalis* (Fig. 2). Similar to previous studies [13, 22], we found that 50 % of wild-type animals fed *E. faecalis* on nematode growth media (NGM) were killed with a median survival (LT_50_) of 9 ± 3 days, post infection. By contrast, 50% of *pbo-1* mutants were killed after an average of 4 ± 1 days of *E. faecalis* feeding, which was significantly shorter than the wild-type animals (*p*<0.005). Exposing wild-type *C. elegans* to RNAi *pbo-1* prior to *E. faecalis* infection also increased susceptibility to *E. faecalis* (Fig. S1). These results indicate that *pbo-1* mutants are more susceptible to killing by *E. faecalis* than the wild type. Because the *pbo-1* mutant causes defects in muscle contraction and defecation, which might slow pathogen clearance, we also tested *pbo-4* mutants. *pbo-4* mutants were not significantly more susceptible than wild type animals with 50 % killed on average 7 ± 2 days by *E. faecalis*. To determine if the increased pathogen sensitivity of *pbo-1* mutants is due to a general defect in lifespan, we grew animals on nonpathogenic *E. coli*. The lifespan of wild type, *pbo-1* and *pbo-4 C. elegans* mutants were similar (median survival was 12-13 days on average) (Fig. 2D). Together, the results show that the *pbo-1* mutant is more sensitive to killing by *E. faecalis* than the wild type, and that this sensitivity is not due to differences in longevity and cannot be entirely explained by defects in posterior body contractions during defecation.

**Fig. 2.**
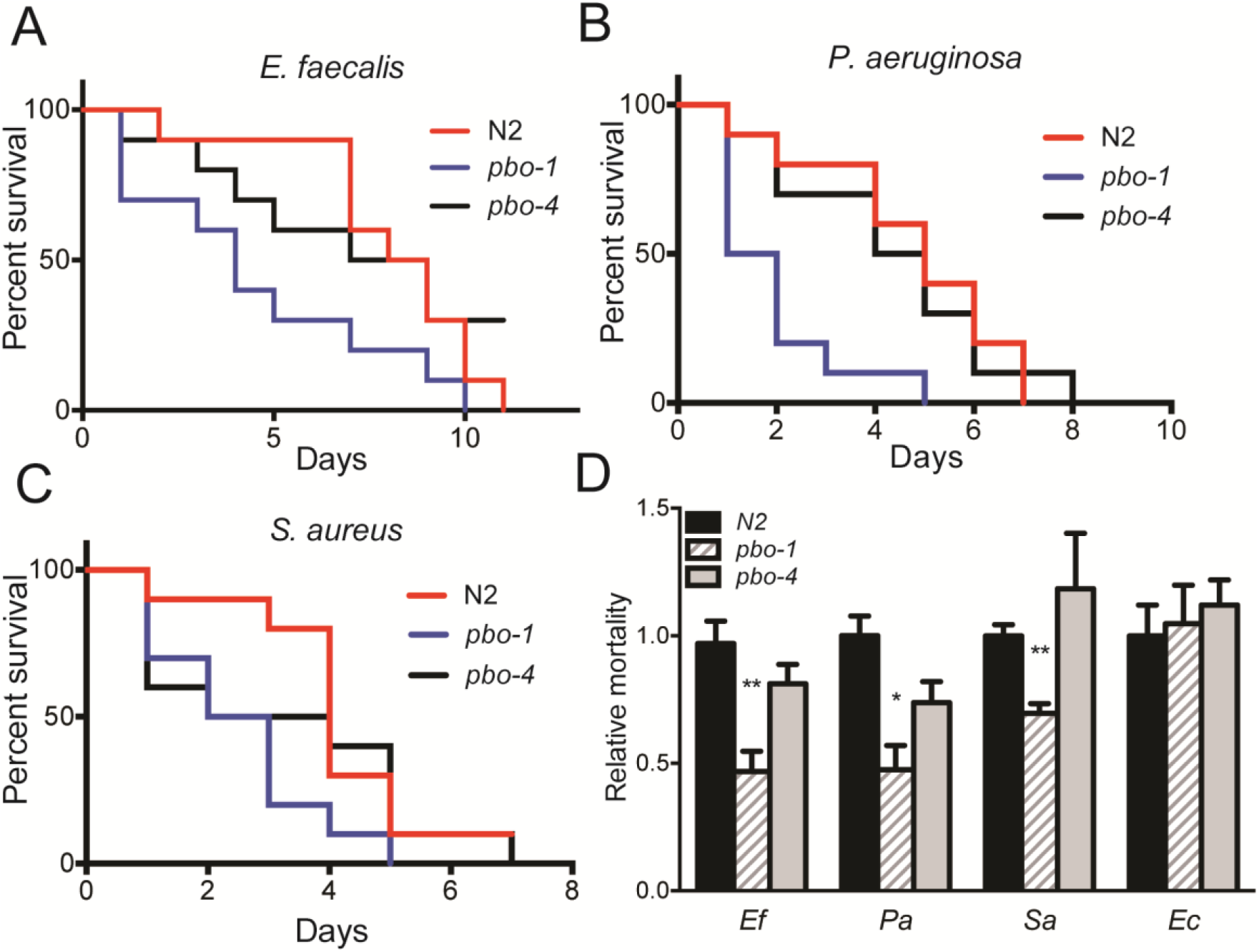
*pbo-1* loss-of-function increases susceptibility to bacterial pathogens. (A-C) Survival of *C. elegans* wild type, *pbo-1, and pbo-4* mutants placed on a lawn of *E. faecalis* (A), *P. aeruginosa* (B), or *S. aureus* (C). In each experiment 10 worms were placed on a lawn with the bacterial pathogen and transferred every two days to a fresh lawn. Representatives from one experiment with 10 worms are shown. In each case, wild type was statistically different from *pbo-1* by log-rank (Mantel-Cox) test. (B) Relative mortality on pathogen (*E. faecalis, P. aeruginosa* or *S. aureus*) or longevity on *E. coli* is shown as a ratio of the LT_50_ (calculated median survival) of the mutant or average wild type *C. elegans* over the LT_50_ of wild-type *C. elegans* for each experiment. Relative mortality was calculated from the average of five (Ec) or three (Pa, Sa and Ec) independent experiments, with 10 animals each. For each pathogen, statistical analysis by student’s *t* test compared with wild type: **, *p<*0.005; *, *p*<0.05.

To determine if the increased susceptibility of *pbo-1* mutants was limited to *E. faecalis*, we tested susceptibility of the *pbo-1* mutant to two other known *C. elegans* pathogens, *Pseudomonas aeruginosa* and *Staphylococcus aureus*. In each case, loss of function in *pbo-1* was associated with increased susceptibility as observed for *E. faecalis* (Fig. 2). For both *P. aeruginosa* and *S. aureus*, survival of the *pbo-4* mutant was not statistically different than wild type (Fig. 2). These results indicate that animals lacking *pbo-1* were more susceptible than wild type to a wide class of pathogens. Together, the results suggest *pbo-1* might be functioning as part of the innate immune system, responsible for protecting the animal from such pathogens.

### Bicarbonate treatment reverses pbo-1 susceptibility

Our results indicate *pbo-1* is important for regulating intestinal pH in *C. elegans*, consistent with studies reported on the *pbo-1* homolog in vertebrates, including humans [23, 24]. To more directly test whether the abnormally acidic intestinal pH contributes to pathogen susceptibility of the *pbo-1* mutant we carried out infections on plates containing sodium bicarbonate (25 mM, pH = 7). This buffer was chosen because chyme, the acidic and partially digested fluid that leaves the stomach of vertebrates, is buffered by the secretion of bile acids from the gall bladder and bicarbonate from the pancreas as it is transported to the small intestine, and bicarbonate is a physiologic buffer for acid in the alimentary canal.

We found that *pbo-1* mutants treated with bicarbonate during *E. faecalis* exposure were less susceptible to killing by the pathogen (median survival was 10 ± 2 on 25 mM bicarbonate, compared with 5 ± 1 days with no bicarbonate) (Fig. 3). Treatment with bicarbonate did not provide wild type or *pbo-4* mutants any additional resistance to *E. faecalis*. Thus, we concluded that the increased susceptibility to pathogens was, at least partially, due to the increased acidity of the alimentary canal of *pbo-1* animals.

**Fig. 3.**
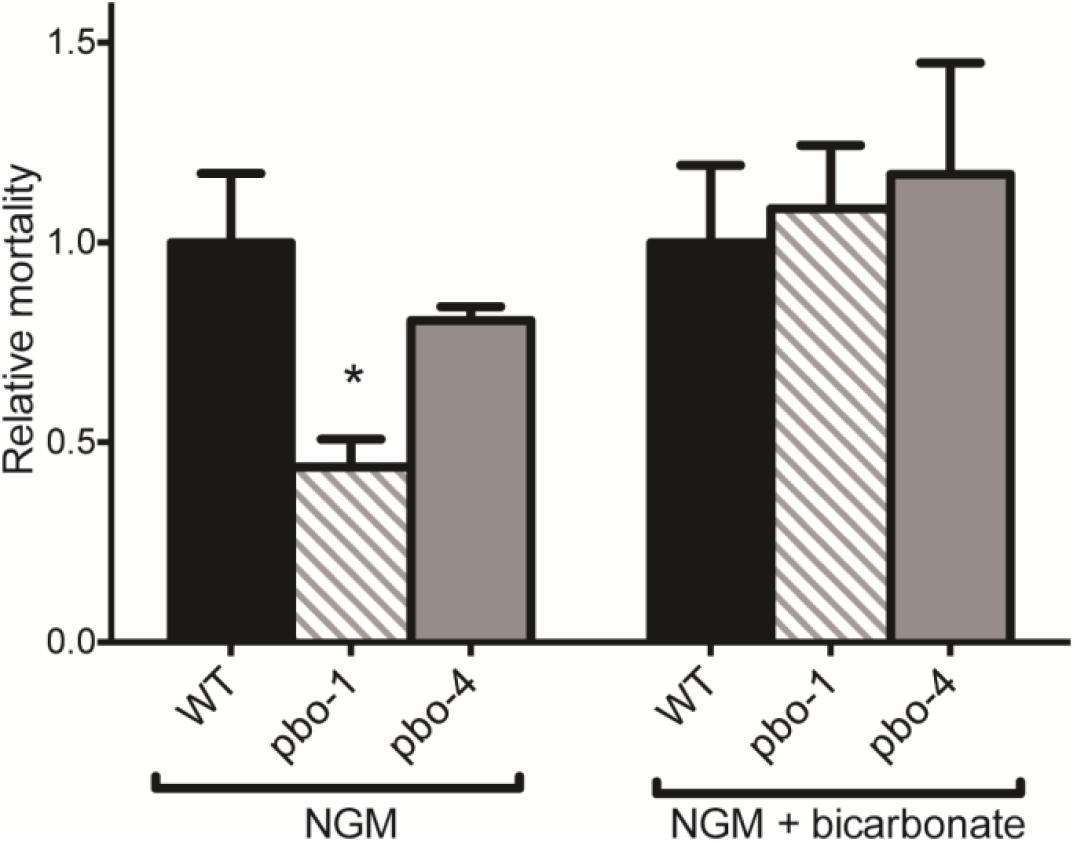
Bicarbonate increases survival of *E. faecalis-fed pbo-1* mutants. Survival of wild-type, *pbo-1* or *pbo-4* mutants fed *E. faecalis* on nematode growth medium (NGM) or NGM with 25 mM bicarbonate. Relative mortality is shown as a ratio of the LT_50_ of mutant or average wild type *C. elegans* over the LT_50_ of wild-type *C. elegans* for each condition. Relative mortality was calculated using the average of two or five independent experiments (for NGM or NGM + bicarbonate, respectively), each with 10 animals. Statistical analysis by student’s *t* test compared with wild type: *, *p*<0.05. WT was not statistically different from either mutant on NGM + bicarbonate.

Because intestinal acidity is linked to pathogen susceptibility, and because removing PBO-4 from *pbo-1* mutants reduced the observed acidity of the alimentary canal (Fig. 1), we tested the susceptibility of *pbo-1; pbo-4* double mutants to *E. faecalis*. These animals were more resistant than the *pbo-1* single mutant, similar to the *pbo-4* single mutant (Fig. S2). Together, our results show that either chemically or genetically reducing acidity in *pbo-1* mutants resulted in increased survival when challenged with pathogens.

### Exposure to E. faecalis affects the pH in the intestine

The ability of bicarbonate treatment to suppress pathogen susceptibility in the *pbo-1* mutant suggested that modulation of intestinal pH might be part of the normal response to pathogens. To test this idea, we compared intestinal acidity of the wild-type *C. elegans* fed nonpathogenic *E. coli* (strain OP50) or a pathogen, either *E. faecalis* or *P. aeruginosa*, using the KR35 dye. We found that animals fed OP50 had the expected dynamic pH wave, with a normal periodicity of ~50 seconds. In contrast, KR35 fluorescence was reduced in animals fed *E. faecalis* or *P. aeruginosa*, indicating the intestinal pH was more neutral in pathogen-fed animals (Fig. 4, Supp. Videos 4 and 5). This suggests that part of the normal response to infection is to neutralize the intestinal pH.

**Fig. 4.**
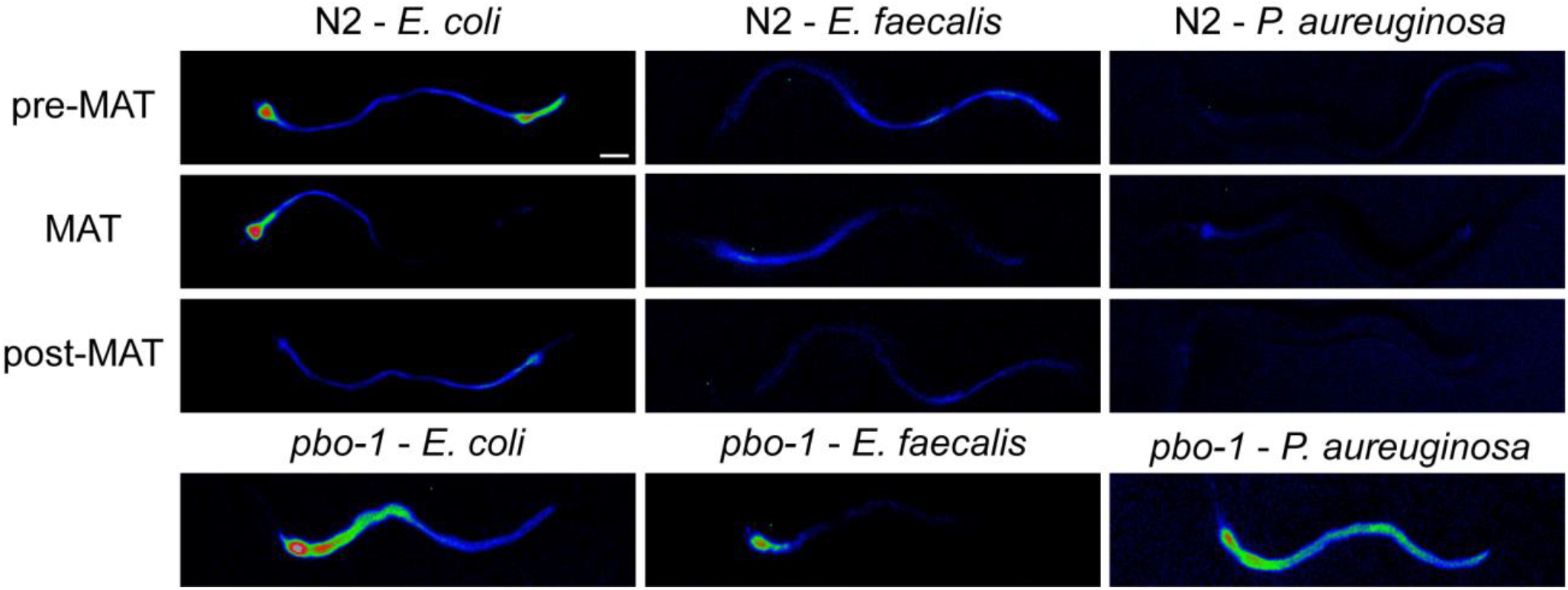
Pathogens alter *C. elegans* intestinal pH. Fluorescence micrographs of *C. elegans* wild type or *pbo-1* mutant fed *E. coli* or pathogen (*P. aeruginosa* or *E. faecalis*) for 1 hour, followed by administration of KR35 (10 M) for 30 min. Time-dependent images during the DMP extracted from video microscopy are shown. Fluorescence is rendered as a heat map of fluorescence intensity with red representing the most intense fluorescence (high acidity), and black the least intense fluorescence (low acidity). The head of the animal is on the left side of each image. (scale bar = 50 μm).

We subsequently tested whether pathogen exposure induced pH changes in *pbo-1* mutants. Similar to wild-type animals, the KR35 fluorescence intensity was reduced in *pbo-1* mutants fed pathogens when compared to *E. coli*-fed mutants. However, in contrast to wild-type animals, *pbo-1* mutant intestinal pH remained acidic on pathogens (Fig. 4), and no pH oscillations were observed (Supp. Videos 6 and 7). Thus PBO-1 is needed to completely neutralize intestinal pH in response to pathogens, but other factors are likely to also be involved in this response.

### pbo-1 mutants exhibit reduced H_2_O_2_ in response to E. faecalis

Work from the Ausbel and Garsin labs have shaped our understanding of the molecular events of pathogenesis, and specifically how *E. faecalis* and *C. elegans* interact during infection. Briefly, recent work has identified that infection of *C. elegans* results in the production of H_2_O_2_ as part of the innate immune system [12]. The BLI-3/Duox and SKPO-1 proteins appear to function in the extracellular space of the intestine to produce H_2_O_2_ and, perhaps convert the H_2_O_2_ into bactericidal compounds. Members of the peroxidase family of enzymes have been established to have pH-dependent activity [25, 26]. Thus, we hypothesized that *pbo-1* or infection-dependent changes in pH might regulate the production of H_2_O_2_.

The Amplex Red assay (see Materials and Methods) can report on the amounts of H_2_O_2_ produced in response to infection. We used this method to quantify H_2_O_2_ produced by wild-type and *pbo-1* mutant animals infected with *E. faecalis* or an *E. coli* control. Consistent with previous results [12], the fluorescent product of the Amplex Red assay was increased in animals infected with *E. faecalis*, as compared with *E. coli*-infected animals (Fig. 5A). However, in the *E. faecalis-*infected animals, the fluorescent product was reduced in *pbo-1* mutants compared with wild type (Fig. 5A). The fluorescent product was similarly reduced in infected animals exposed to RNAi *pbo-1* (Fig. S3). Because bacteria can produce H_2_O_2_ in addition to *C. elegans*, to show that the changes in H_2_O_2_ were due to the worm, and not *E. faecalis*, we used an *E. faecalis* strain deficient for production of H_2_O_2_ (*menB* mutant). As previously reported, animals fed the *E. faecalis menB* mutant had levels of Amplex Red fluorescent product that were similar to that of wild-type *E. faecalis-*fed animals (Fig. S4). These results support that *pbo-1* mutants are defective for production of H_2_O_2._ and are consistent with the idea that defects in production of H_2_O_2_ in *pbo-1* mutants can account for the increased susceptibility to pathogens.

**Fig. 5.**
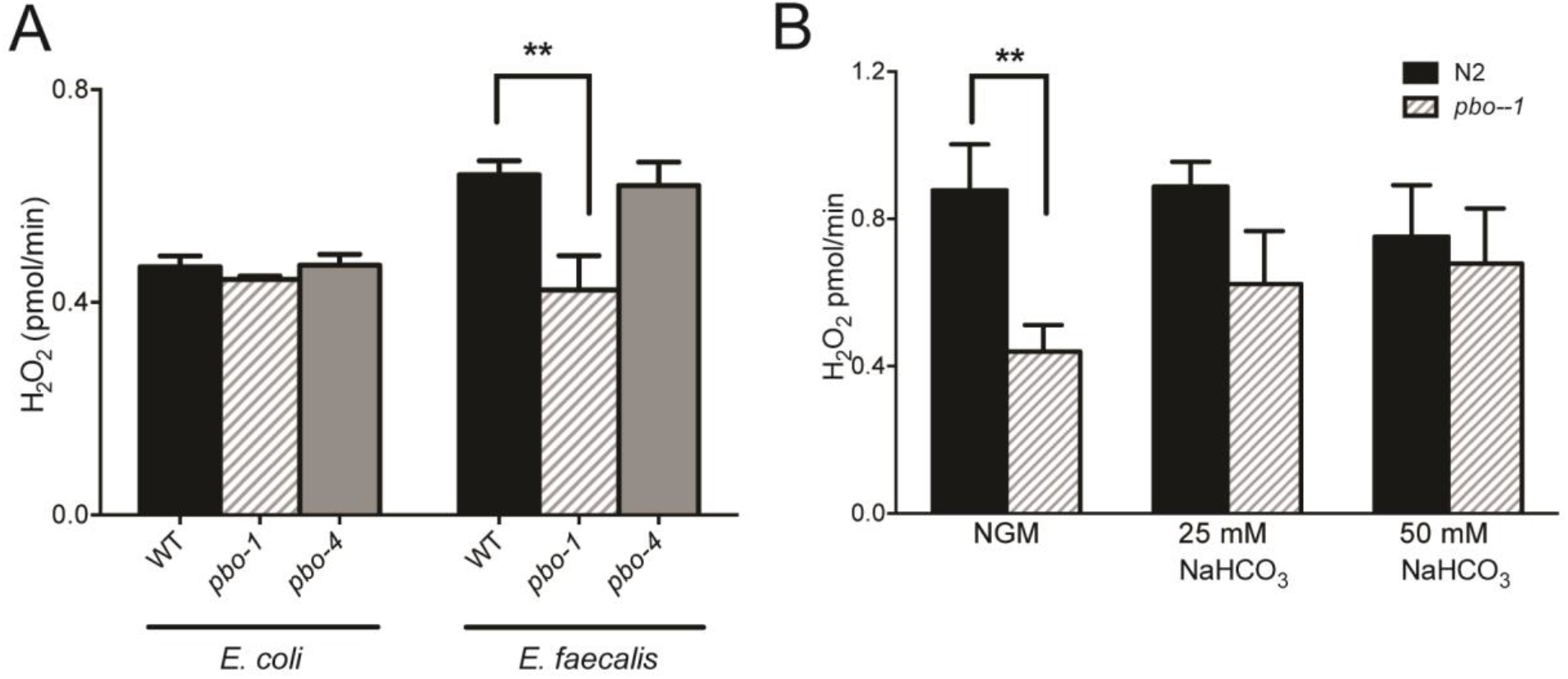
Amplex Red measurements of H_2_O_2_. (A) PBO-1, but not PBO-4, is important for H_2_O_2_ production on in response to *E. faecalis*. (B) In the presence of *E. faecalis*, H_2_O_2_ is restored to a *pbo-1* mutant by adding bicarbonate (NaHCO_3_) to the nematode growth medium. Results are of three independent experiments. Error bars represent standard deviation. **, statistically different by student’s t-test (*p* < 0.006). WT and *pbo-1* were not statistically different on NGM + 25 or 50 mM bicarbonate.

To test whether H_2_O_2_ production is regulated by changes in pH, we neutralized the intestinal pH of *pbo-1* mutant animals with bicarbonate during infection, then used the Amplex Red assay to quantify H2O_2_. Our results showed that the fluorescent product in bicarbonate-treated *pbo-1* mutants was not statistically different from that of wild type (Fig. 5B). Thus, bicarbonate reverts the defect in H_2_O_2_ production in *pbo-1* mutants. These results support the conclusion that the reduction of H_2_O_2_ production in the *pbo-1* mutant is pH dependent.

Because *pbo-4* mutants have an opposite effect on pH in the intestine from that of *pbo-1* mutants, we also tested *pbo-4* mutant for Amplex Red activity. Interestingly, the *pbo-4* mutant showed an intermediate Amplex Red phenotype between the wild type and the *pbo-1* mutant (Fig. 5A). Although *pbo-4* mutants have less acidic pH than wild type, it is possible that defects in MAT and the acidic wave in the *pbo-4* mutant might cause pH-dependent effects on the enzymes that produce H_2_O_2_ similar to, but not as severe as, the *pbo-1* mutant. It is of note that *pbo-4* mutants were slightly, though not statistically, more susceptible to pathogens than wild type (Fig. 2), consistent with the idea that pathogen susceptibility and H_2_O_2_ production are correlated. Together, the results support the idea that a normally functioning MAT and acidic wave are important for *C. elegans* to defend against pathogens via production of H_2_O_2_.

## Discussion

We are interested in characterizing the dynamics of changes in acidity of the *C. elegans* intestine and understanding how these changes impact worm physiology. To characterize intestinal acidity, we previously developed the acid-activated fluorophore KR35 [18], which affords an advantage over other pH-sensitive dyes, *e.g*. Oregon Green, that become non-fluorescent under acidic conditions. KR35 has provided a unique view into the dynamic pH changes that occur in the *C. elegans* intestine when animals are maintained on non-pathogenic *E. coli* [18]. A major finding of the current study is that when we used KR35 to characterize the intestinal pH of animals grown on pathogens, we discovered that the intestine became more alkaline along the length of the lumen, suggesting changes in pH are part of the normal response to infection.

This begs the question as to whether changing the pH is a by-product of the stress response to pathogens or is a critical function itself of fighting pathogens. We addressed this question using a mutant disrupted in the EF-hand protein, PBO-1. PBO-1 is a critical regulator of ion channels that contribute to the pH of the intestine. Our finding that *pbo-1* mutants are more acidic than wild type in the anterior region of the intestine (Fig. 1), and that *pbo-1* mutants are also more susceptible to pathogens (Fig. 2) supports the idea that regulation of pH in the intestine is an important component of pathogen defense. Critically, exposing *pbo-1* mutants to bicarbonate, which neutralizes intestinal pH, reverted susceptibility (Fig. 3). Although PBO-1 is also involved in defecation [14], the wild-type susceptibly of *pbo-4* mutants (also defecation defective) and *pbo-1, pbo-4* double mutants suggested that normal defecation was not required for the animal to respond and clear the pathogen (Fig. 2). These results support the idea that intestinal pH is linked to anti-pathogen activity, independent of its role in defecation.

The pH of extracellular spaces has been shown in other contexts to be important for fighting pathogens. In a porcine CF model, the airway surface liquid (ASL) is more acidic than normal animals, and the ASL shows reduced killing of bacterial pathogens [27, 28]. In that model, the authors show that they can also use bicarbonate to simply, and rapidly, restore the antibacterial properties of the ASL [28]. The antimicrobial activity was proposed to be due to pH-modulated antimicrobial peptides (AMPs). We further found that in *C. elegans* with acidic intestines (*pbo-1* mutant), pH modulated the production of hydrogen peroxide (H_2_O_2_) (Fig. 5). H_2_O_2_ has been shown to be an important signaling molecule in physiological contexts such as wound healing and infection. It may be involved in reporting on the presence of pathogens or in the formation of bactericidal agents that kill microbial invaders. Whether this represents increased production or stability remains an open question. Together, these results support that pH modulation of innate immune responses might be conserved across animal classes and systems.

pH also modulates other immune functions. In *in vitro* studies, pH affects the activity of dendritic cells [29], neutrophils [30-32], and the complement system [33]. In patients, lower pH is often associated with reduced immune function [34], and reduced production of bicarbonate is linked with high inflammation [35]. We have shown that in *C. elegans*, regulation of pH appears to be tied to the PBO-1/PBO-4 pathway. These pathways have previously been studied in the context of their role in regulating defecation. PBO-1 modulates PBO-4 [20], an Na^+^/H^+^ ion exchange pump that normally exports protons from the basolateral membrane of the intestine to adjacent muscles, causing stimulation of the contraction of the posterior body wall muscles prior to defecation [17]. Our results show that PBO-1/PBO-4 might also help to clear pathogens through pH-dependent changes in H_2_O_2_ production. It is reasonable to propose that ingestion of pathogens triggers a shift from defecation (muscle contraction) to pathogen defense (production of H_2_O_2_) in *C. elegans*. The PBO-1 and PBO-4 proteins are conserved in mammalian cells [15, 16] and it will be interesting to determine whether these proteins have a similar role in regulating pH and immune function in higher species.

## Materials and Methods

### Strains and culture conditions

*C. elegans* and bacterial strains used in this study are listed in Table 1. All *C. elegans* strains were maintained on nematode growth medium (NGM) at 20°C as previously described [36]) unless otherwise noted. The reference strain for all alleles used and generated was N2 (var. Bristol). All bacterial strains were grown at 37ºC. *E. coli* and *P. aeruginosa* bacterial strains were grown in low-salt Luria Bertani medium (10 g Bacto-tryptone, 5 g yeast extract, 5 g NaCl), and *E. faecalis* and *S. aureus* were grown on Brain Heart Infusion, Porcine (BHI, from BD). For *E. faecalis*, broth cultures were incubated without shaking. All other bacterial strains were grown with shaking. For selection and expression of RNAi clones, ampicillin (50 μg/mL), tetracycline (15 μg/mL) and isopropyl β-D-1-thiogalactopyranoside (IPTG) (1 mM) were added to the growth medium.

### *C. elegans* survival and longevity assays

For survival assays, NGM plates were spread with 10 μL stationary-phase bacteria (*E. faecalis, S. aureus*, or *P. aeruginosa* grown to an optical density at 600 nm [OD_600_] ~1), and the plates were incubated overnight at 37 °C. Plates were acclimated to room temperature for ~1 hour prior to seeding the plates with *C. elegans* (10 late larval (L4) staged worms per plate). Worms were scored for survival each day and transferred to new NGM plates with bacterial lawns every two days. Where indicated in the text, NGM was supplemented with 25 mM sodium bicarbonate buffer (pH 7). Longevity assays on *E. coli* were conducted as described for survival assays, except NGM plates were spread with 40 μL *E. coli* OP50 and incubated overnight at 20 °C prior to seeding with *C. elegans*.

### RNAi

RNAi was performed by exposing L1-L4-stage larvae to a lawn of *E. coli* HT115-expressing dsRNA to target genes on NGM plates for 3 days prior to performing survival or Amplex Red experiments. RNAi clone III-7I12 (*pbo-*1) was obtained from the *C. elegans* library (Fraser et al. 2000; Kamath et al. 2003).

### Amplex Red assay for H_2_O_2_ measurements

The Amplex Red hydrogen peroxide/peroxidase kit (Invitrogen Molecular Probes, Eugene, OR) was used to measure pathogen-stimulated hydrogen peroxide release by *C. elegans* as previously described (Chavez et al., 2007, 2009). Briefly, L4 worms were exposed to *E. faecalis* for 24 h, and the fluorescence of 10 worms per well was measured after 30 min incubation in the dark with Amplex Red/peroxidase solution (540/590 excitation and emission, respectively). The quantity of H_2_O_2_ was calculated using an etalon curve.

### Image analysis

KR35 imaging was carried out as described (Bender et al., 2013), with minor modifications. For non-pathogen videos, animals were reared on NGM plates with OP50, and imaged on plates containing OP50. For pathogen-treated animals, animals were reared on OP50, then transferred to NGM plates containing either *E. faecalis* or *P. aeruginosa*, as indicated. Prior to acquisition of videos, young adults were transferred to NGM plates supplemented with 10 M KR35, for 30 minutes, and then imaged on NGM plates with the indicated bacterial strain. Images were acquired on a Leica M165FC microscope using a Leica DFC3000G CCD Camera via the Leica Application Suite software (v 4.40) (Leica Microsystems (Switzerland) Limited). Single images and image sequences were acquired at 5x zoom, with 10x gain, and sequences at 10 frames per second. Illumination was via a Leica Kubler Codix source equipped with an Osram HXP 120W lamp. Images were opened in Fiji (ImageJ), converted to 8-bit, scaled from 0-255 to a dynamic range of 10-120, and a rainbow RGB look up table applied. Movies were converted to AVI using FIJI using an approximate 30 s clip that corresponded to ~10-15 seconds before and after an MAT (where one occurred).

## Acknowledgements

We would like to acknowledge Ajai Dandekar for helpful comments on this manuscript. We would also like to thank Danielle Garsin for the detailed Amplex red protocol and helpful comments on this project. This work was supported by the National Institutes of Health under award number P20 GM103638 (to JRC), P20 GM113117 (to BDA), P20 GM103638 (to BRP), and a K-INBRE postdoctoral fellowship award to SB (P20 GM103418).

## Author Contributions

Conceptualization: BRP, JRC and BDA.

Investigation: SB, AB and BDA

Supervision: JRC and BDA

Writing: SB, JRC and BDA

Review & editing: SB, JRC, BDA and BRP

---

**Fig. S1.**
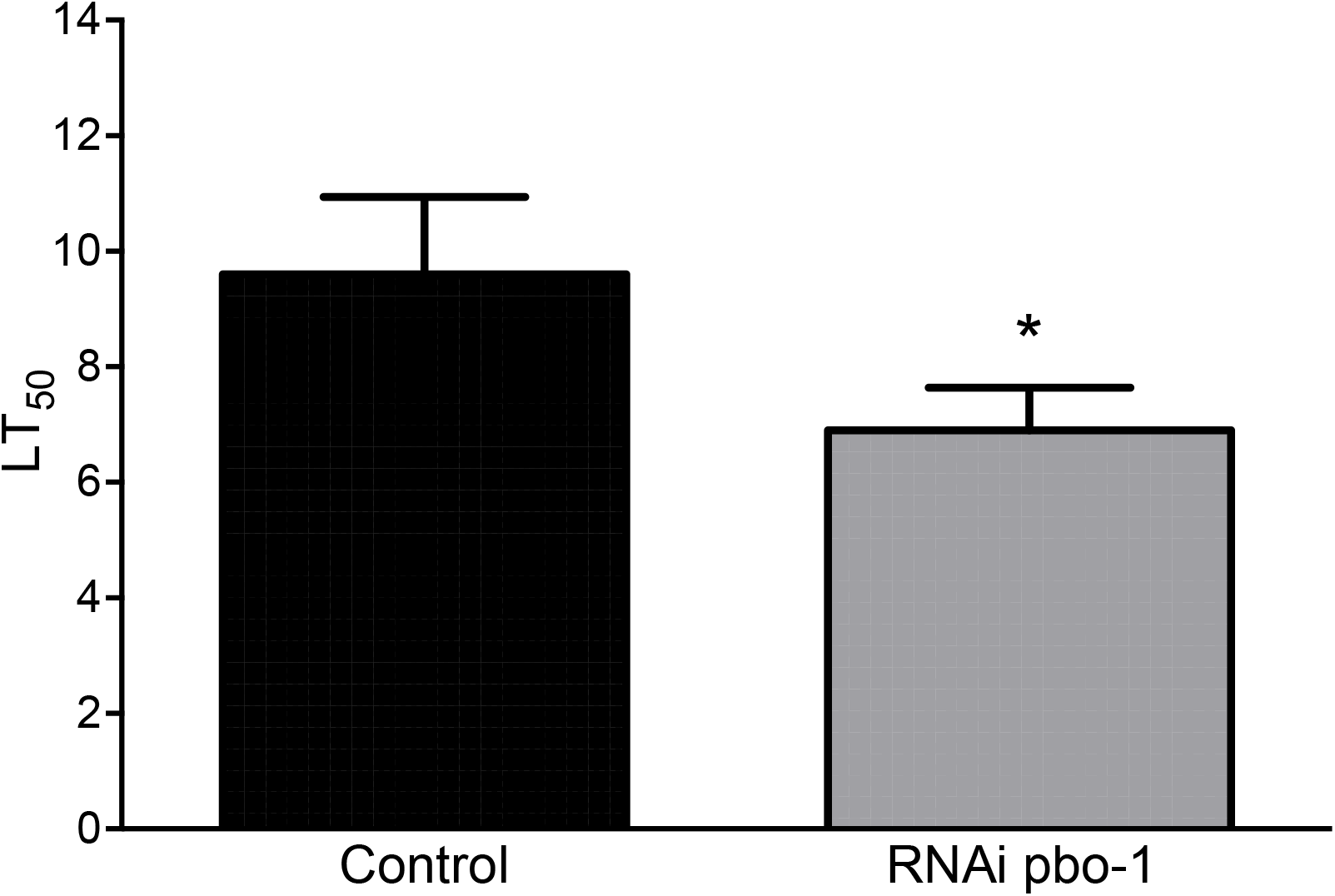
Exposure to RNAi *pbo-1* increases susceptibility to *E. faecalis*. RNAi was performed by exposing L1-L4-stage larvae to a lawn of *E. coli* HT115-expressing *pbo-1* dsRNA (clone III-7I12, obtained from the *C. elegans* library (Fraser et al. 2000; Kamath et al. 2003)) for 3 days prior to transferring to *E. faecalis* for survival experiments. Control, identically treated *C. elegans* grown on *E. coli* OP50. In each experiment, 10 worms were placed on a lawn with *E. faecalis* and transferred every two days to a fresh lawn. Results are of 5 independent experiments performed on two separate days. Error bars represent the standard deviation. *, *p* < 0.005.

**Fig. S2.**
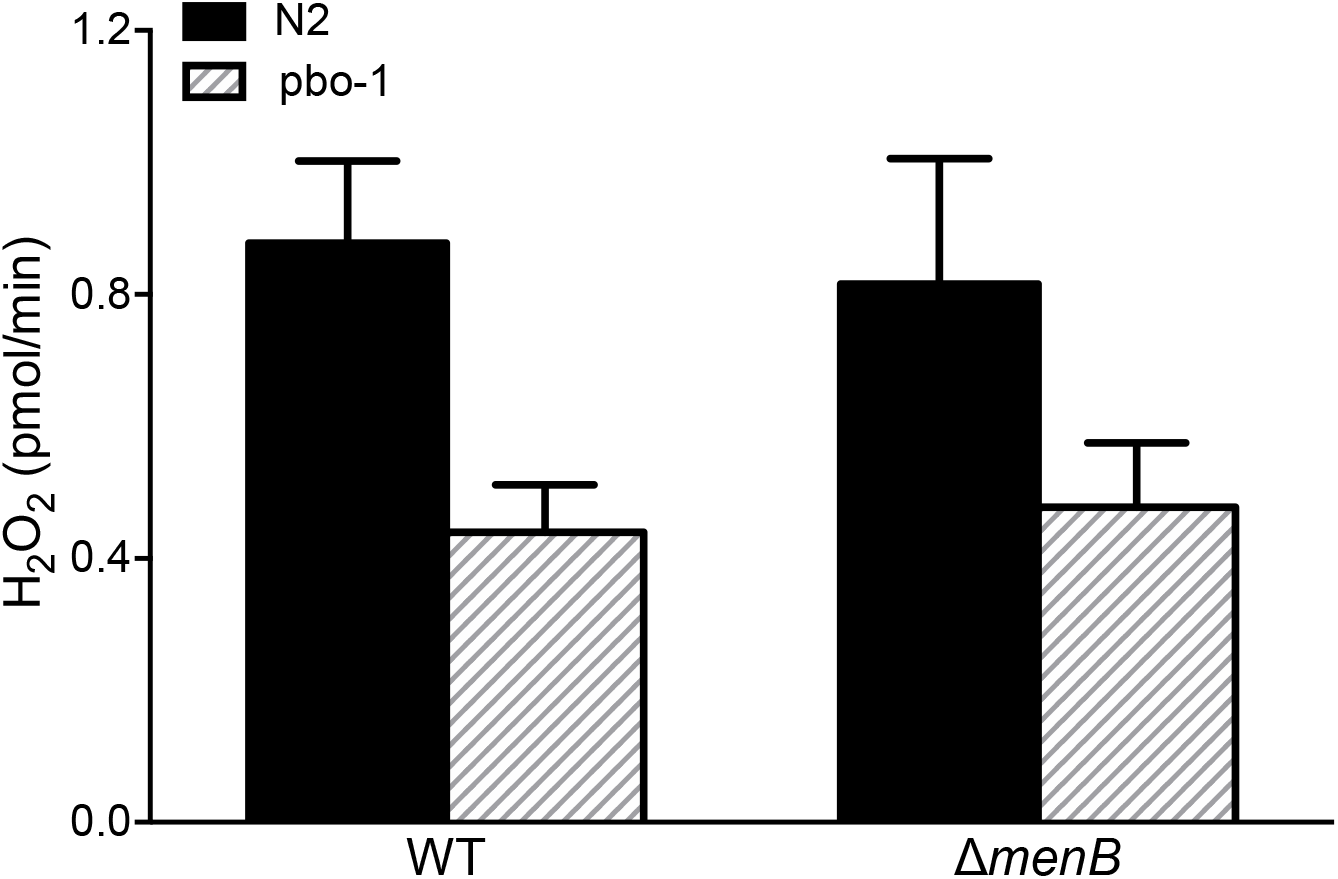
Amplex Red H_2_O_2_ measurements of the H_2_O_2_-deficient *E. faecalis* strain (∆*menB*). Results are the averages of at least four independent experiments performed on two separate days. Error bars represent standard deviation. H_2_O_2_ production is not statistically different between WT and ∆*menB* for both N2 or *pbo-1 C. elegans* (*p*>0.1).

**Fig. S3.**
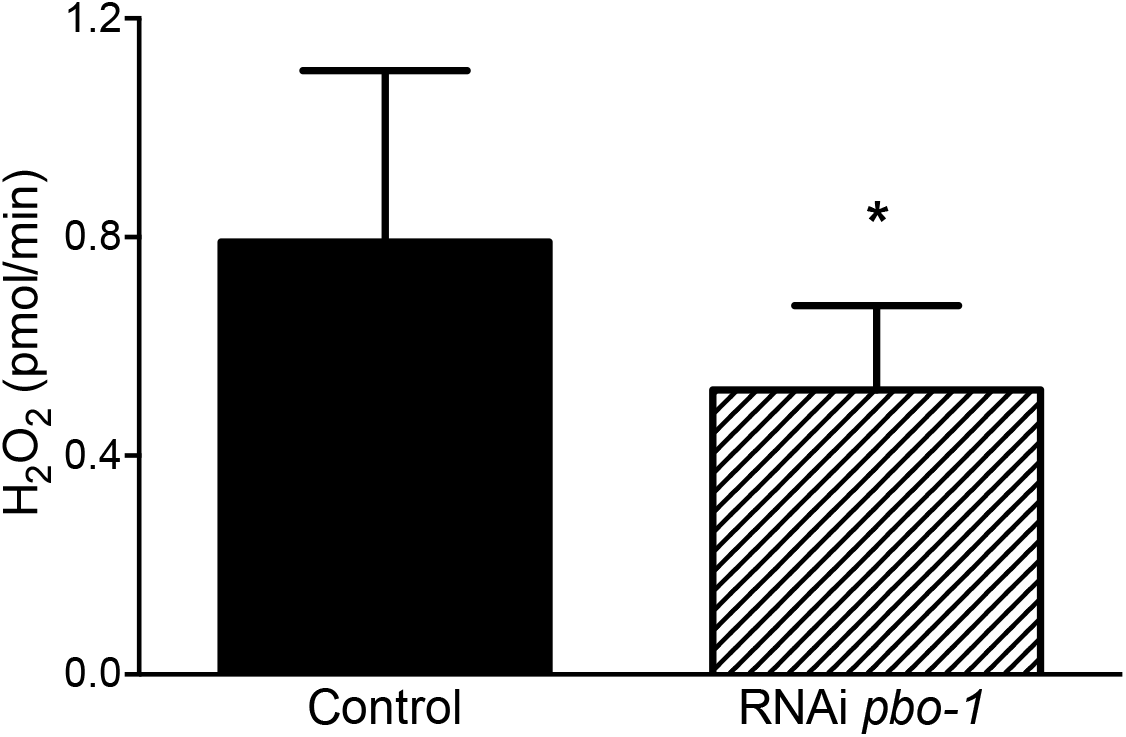
Amplex Red H_2_O_2_ measurements of *C. elegans* exposed to RNAi *pbo-1*. are the averages of four independent experiments performed on three separate days using Amplex Red. Error bars represent standard deviation. *, *p*<0.05 by student’s paired t-test.

**Fig. S4.**
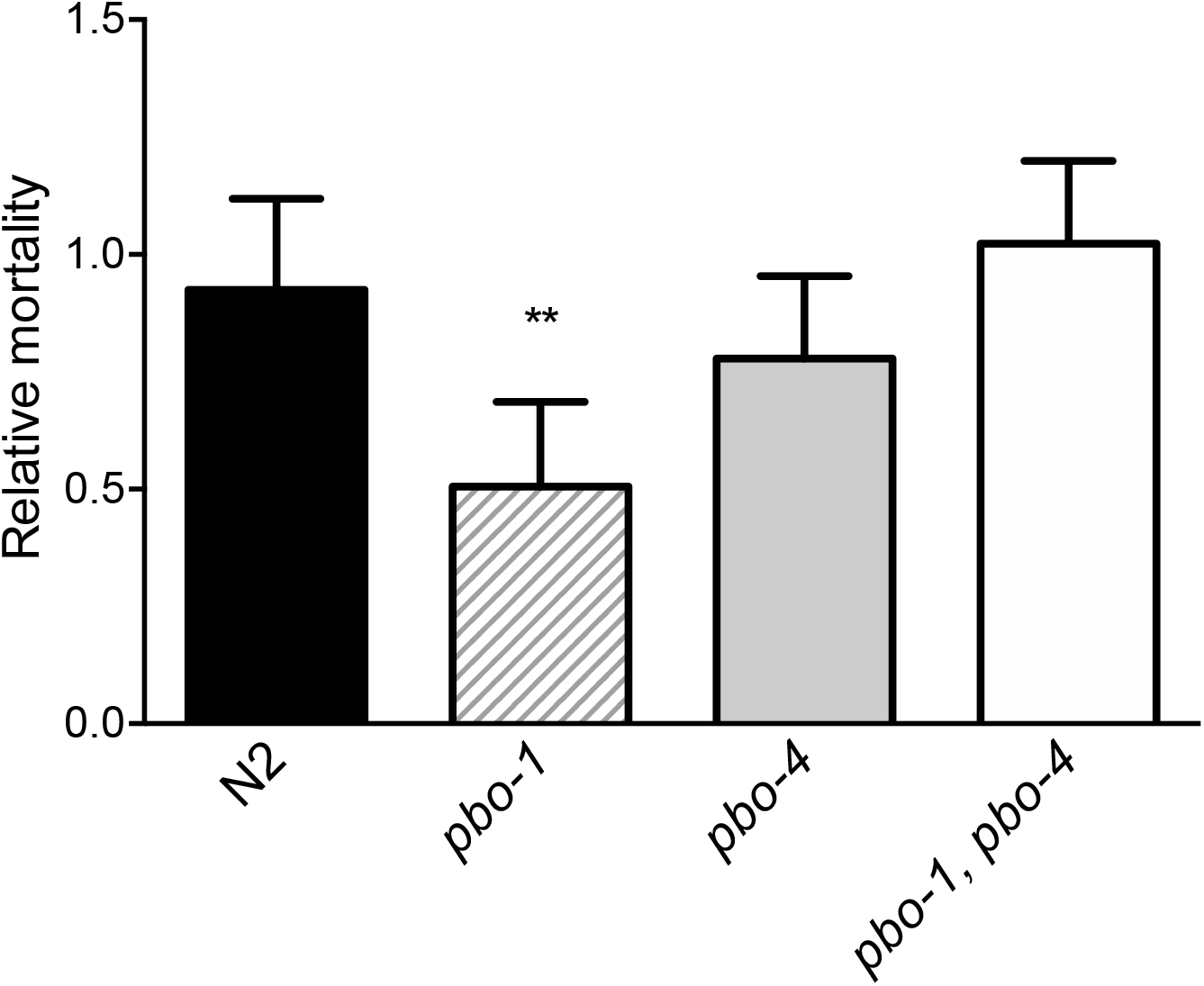
Survival on *E. faecalis is similar for pbo-1* and *pbo-1, pbo-4* double mutants. Relative mortality on pathogen *E. faecalis* is shown as a ratio of the LT_50_ (calculated median survival) of wild type or mutant *C. elegans* over the LT_50_ of wild-type *C. elegans* for each experiment. Relative mortality was calculated from the average of four independent experiments, with 10 animals each. For each pathogen, statistical analysis by *t* test compared with wild type: **, *p<*0.005. Data are also partially represented in Fig. 2D.

**Table S1.** *C. elegans* and bacterial strains used in this study.

| Strain | Relevant properties | Reference or source |
| --- | --- | --- |
| <u>*Caenorhabditis elegans*</u> |  |  |
| N2 | var. Bristol wild-type | [1] |
| RB793 | *pbo-4(ok583)* | [2] |
| JT7 | *pbo-7(sa7)* | [3] |
| EVL1708 | *pbo-1(sa7); pbo-4(ok583)* | This study |
| EVL949 | *pbo-4(ok583); lhEx290 [Ppbo-4::PBO-4::GFP]* | [4] |
| EVL1678 | *pbo-1(sa7); lhEx290* | This study |
| <u>Bacterial strains</u> |  |  |
| *E. coli* OP50 | uracil-requiring mutant | [1] |
| *E. coli* HT115 | dsRNA expression strain for RNAi experiments | [5];<br>[6] |
| *E. faecalis* OG1RF | Derived from clinical strain OG1; carries<br>spontaneous mutations to rifampicin and<br>fusidic acid resistance | [7] |
| *P. aeruginosa* PA14 | Clinical isolate from a human burn patient | [8] |
| *S. aureus* Newman | Early methicillin sensitive isolate from<br>secondary infection in a patient with tubercular<br>osteomyelitis (Sm sensitive) | [9] |

